# Establishment of a murine resident dermal macrophage cell line

**DOI:** 10.64898/2026.05.15.725415

**Authors:** Stephanie L. Brandt, Anderson Sá-Nunes, Ana C.G. Salina, Allison Judge, Dante Reyna, Amondrea Blackman, Nathan Klopfenstein, C. Henrique Serezani

## Abstract

Resident dermal macrophages (DMs) play essential roles in maintaining skin homeostasis and initiating inflammatory responses during tissue injury and against infectious agents. However, studies of their cellular mechanisms have been limited by their low abundance in steady-state skin and by technical challenges in isolating resident DMs. Here, we describe the generation and characterization of a novel DM cell line, termed SB89. F4/80^+^ skin-resident DMs were sorted and immortalized using J2 retroviral transduction. SB89 cells display a stable, homogeneous macrophage phenotype and distinct surface markers compared with Langerhans cells and alveolar macrophages. Functionally, SB89 cells efficiently phagocytose methicillin-resistant *Staphylococcus aureus* (MRSA), *Escherichia coli*, zymosan particles, and apoptotic cells, and effectively kill MRSA. Importantly, SB89 cells respond to LPS, as evidenced by production of IL-6, TNF, and IL-10, and by MRSA-induced production of inflammatory cytokines, chemokines, and eicosanoids. RNA-seq and gene ontology analyses revealed that SB89 cells elicit stronger responses in innate immunity, cell signaling, and epigenetic regulation than immortalized bone marrow-derived macrophages. SB89 cells are genetically tractable, amenable to gene silencing via RNAi and gene introduction via plasmid transfection. Overall, SB89 cells provide a renewable, dermis-imprinted macrophage model that preserves key functional and transcriptional features of resident DMs while reducing reliance on primary cells and animal models. This cell line represents a powerful platform for mechanistic, genetic, and translational studies in skin immunobiology.

## INTRODUCTION

The skin is a highly specialized organ that serves as a critical barrier between the body and the external environment, providing both mechanical and biological protection against a wide range of threats. Structurally, it is constituted of two main layers: the epidermis, which is mainly composed of keratinocytes; and the dermis, a connective tissue network rich in fibroblasts, collagen, and elastin fibers that confer strength and elasticity. The dermis also contains capillary and lymphatic vessels that support immune cell trafficking, as well as hair follicles, sebaceous glands, sweat glands, and nerve endings^1^.

The skin provides a unique platform for macrophage research because of its distinct anatomical organization and specialized immune cell populations distributed throughout the epidermis and dermis. The epidermis contains Langerhans cells (LCs), which are antigen-presenting cells (APCs) and share a common origin and renewal process with embryonically-derived macrophages^2^, whereas the dermis harbors tissue-resident macrophages (large, vacuolar phagocytic cells) and dermal dendritic cells (also APCs) ^1, 2^. Additional populations of macrophages and T cells are found within the hypodermis. These cell populations can be identified based on characteristic marker expression profiles. Langerhans cells express CD1a, CD11b, CD11c, F4/80, CD207 (langerin), and MHC class II molecules; dermal dendritic cells (DCs) display CD1a^low^, CD1c, CD11b^high^, CD206, CD209, and MHC class II; dermal macrophages express markers including CD11b, F4/80, CD163, factor XIIIa, CD16, CD32, and CD64^1, 2^.

Skin-resident macrophages are essential for maintaining tissue homeostasis, orchestrating immune responses, and promoting skin repair and regeneration^3^. Their significance lies in their diverse and remarkable functional plasticity within the skin’s microenvironment, which serves both as a physical barrier and a highly active immunological interface between the external environment and internal systems. These macrophages continuously patrol the dermis, acting as sentinels that detect pathogens, tissue damage, and abnormal cellular activity^3^. They are crucial for maintaining tissue homeostasis by phagocytosing apoptotic cells, cellular debris, and foreign particles, thereby preventing harmful accumulation of material and preserving tissue integrity^4, 5, 6, 7^.

Interest in dermal macrophages has increased considerably due to their involvement in skin-associated disorders, including psoriasis, atopic dermatitis, chronic wounds, and cutaneous infections, as well as in tumor progression within the skin microenvironment^4, 5, 6, 7^. Despite this growing interest, the field faces significant challenges due to the lack of accessible, physiologically relevant *in vitro* models. Although primary dermal macrophages provide valuable biological insights, their use is restricted by their limited lifespan in culture and the labor-intensive isolation procedures required to obtain them from skin biopsies. In contrast, commonly used immortalized macrophage cell lines are typically derived from non-cutaneous tissues (e.g., bone marrow, peritoneal cavity, or lung) and, therefore, lack the tissue-specific characteristics of dermal macrophages. This gap in model systems limits the ability to study the unique biology of dermal macrophages and their interactions within the skin microenvironment.

Creating a stable dermal-specific macrophage cell line that closely mirrors primary dermal macrophages in both characteristics and functions is essential to addressing these challenges. Such a model would enable long-term, consistent investigation of macrophage roles in the skin, including responses to inflammatory triggers, interactions with skin-resident cells, and involvement in skin diseases. In addition, it would also support high-throughput drug screening and mechanistic studies targeting skin immune responses. Recent research indicates that immortalizing macrophage populations may be a viable solution. This can be achieved in specific mouse strains using the J2 retroviral infection system, which has been successfully applied to fetal liver macrophages, splenic macrophages, microglia, and bone marrow-derived macrophages (BMDM)^8, 9, 10^. Importantly, immortalized murine macrophage lines generated using this approach retain typical macrophage functions and effectively mimic primary cells^10, 11,12^. Furthermore, these cell lines can be genetically engineered to investigate key immunological and metabolic pathways in greater detail^12^.

Research on skin inflammatory responses and immunology is limited to *in vivo* models, making mechanistic studies difficult. Here, we established a continuous line of murine skin-resident DMs termed SB89. This cell line highlights the unique hallmarks of DMs and supports phenotypic and functional characterization. These cells are also well-suited to generating reporter cell lines by introducing modified proteins via transfection. This work lays the foundation for future research into macrophage-mediated skin pathologies and therapeutic interventions targeting dermal macrophage functions.

## RESULTS

### Generation and phenotypic characterization of SB89

Skin biopsies were collected, and single-cell suspensions were FACS-sorted to isolate F4/80^+^ cells. The sorted cells were cultured in the presence of M-CSF plus with 50% AMJ2-C11-expressing J2 retrovirus supernatant. Proliferating cells were observed on day 15 post-challenge and were subsequently maintained in culture with progressively decreasing concentrations of M-CSF. After culturing without exogenous M-CSF, limiting dilution was performed to select five proliferative single-cell clones (Fig. 1A). In suspension, SB89 cells display a rounded morphology, and upon adherence, they tend to elongate and spread across the culture surface. After 3-4 days of culture, the cells acquire a morphologically homogeneous appearance. During the initial stages after seeding, before complete adherence, SB89 cells exhibit a predominantly rounded morphology (Fig. 1B).

**Fig. 1.**
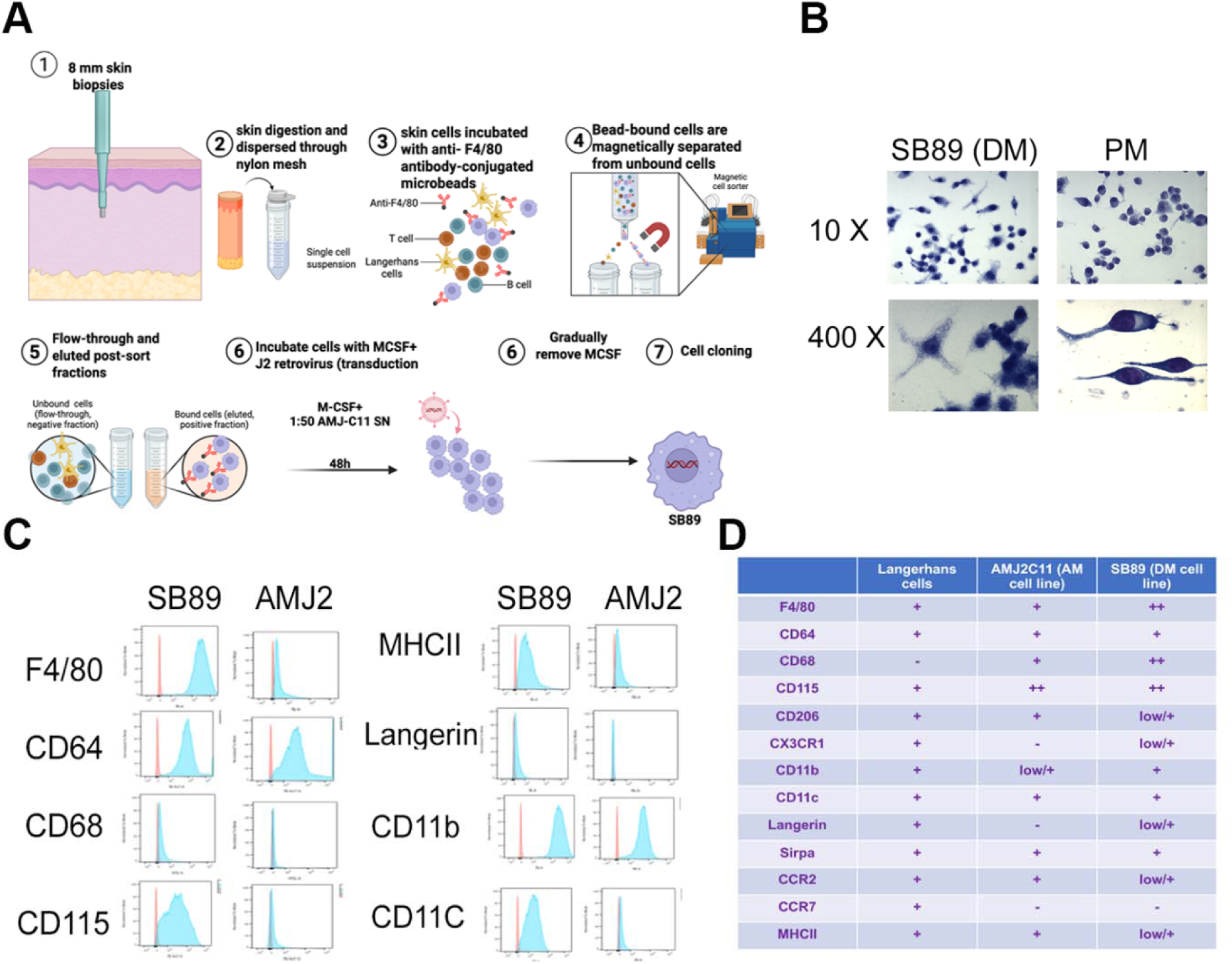
Isolation and characterization of the SB89 dermal macrophage cell line. A) Skin biopsies were collected; skin macrophages were isolated by negative selection and immortalized with J2 retroviruses. **B)** SB89 cells and primary peritoneal macrophages were stained with Diff-Quick as described in the Materials and Methods section. **C)** Expression of macrophage markers in AMJ2-C11 cells was assessed by flow cytometry. **D)** Relative abundance of markers among primary Langerhans cells, AMJ2-C11, and SB89 cells was determined by flow cytometric analysis. Data are representative of at least 4 independent experiments.

SB89, an alveolar macrophage (AMJ2-C11, used as a source of J2 retrovirus^8, 13^), and LCs were each stained with antibodies targeting macrophage and LC markers, then subjected to a series of flow cytometry analyses (Fig. 1C and D). Importantly, LCs were characterized based on the expression of Langerin and CCR7. These cell lines express similar levels of F4/80, CD64, CD11c, SIRPa, and MHC class II molecules. In contrast, DMCL exhibited distinct expression patterns for CD68, CCR2, CX3CR1, and MHC class II compared with AMs and LCs (Table 1). Collectively, our results show that SB89 cells are a homogeneous population (confirming their clonal nature) that maintain skin phagocyte markers and therefore can be used in *in vitro* studies.

### SB89 cells efficiently phagocytose and kill pathogens

To determine whether SB89 cells are suitable for functional immune studies, we assessed their antimicrobial effector functions and their ability to produce inflammatory and anti-inflammatory mediators in response to stimulation. SB89 cells effectively phagocytosed all tested targets (heat-killed methicillin-resistant *Staphylococcus aureus* [HK-MRSA], *Escherichia coli,* and zymosan particles), as shown in Fig. 2A. Additionally, under similar conditions, SB89 cells engulfed significantly more microbial particles than primary peritoneal macrophages (PM) and the AM cell line (Fig. 2B). SB89 cells also efficiently ingested apoptotic cells (Fig. 2C) and effectively killed MRSA (Fig. 2D). Next, we examined whether lipid mediators known to influence macrophage behavior, namely prostaglandin E_2_ (PGE_2_) and leukotriene B4 (LTB_4_)^14, 15, 16, 17, 18^. We found that phagocytosis and bacterial killing by SB89 cells were inhibited by PGE_2_ and enhanced by LTB_4_ (Fig. 2D and E). Together, these results demonstrate that SB89 cells possess robust microbicidal capacity and constitute an appropriate model for experiments investigating how mediators affect macrophage function.

**Fig. 2.**
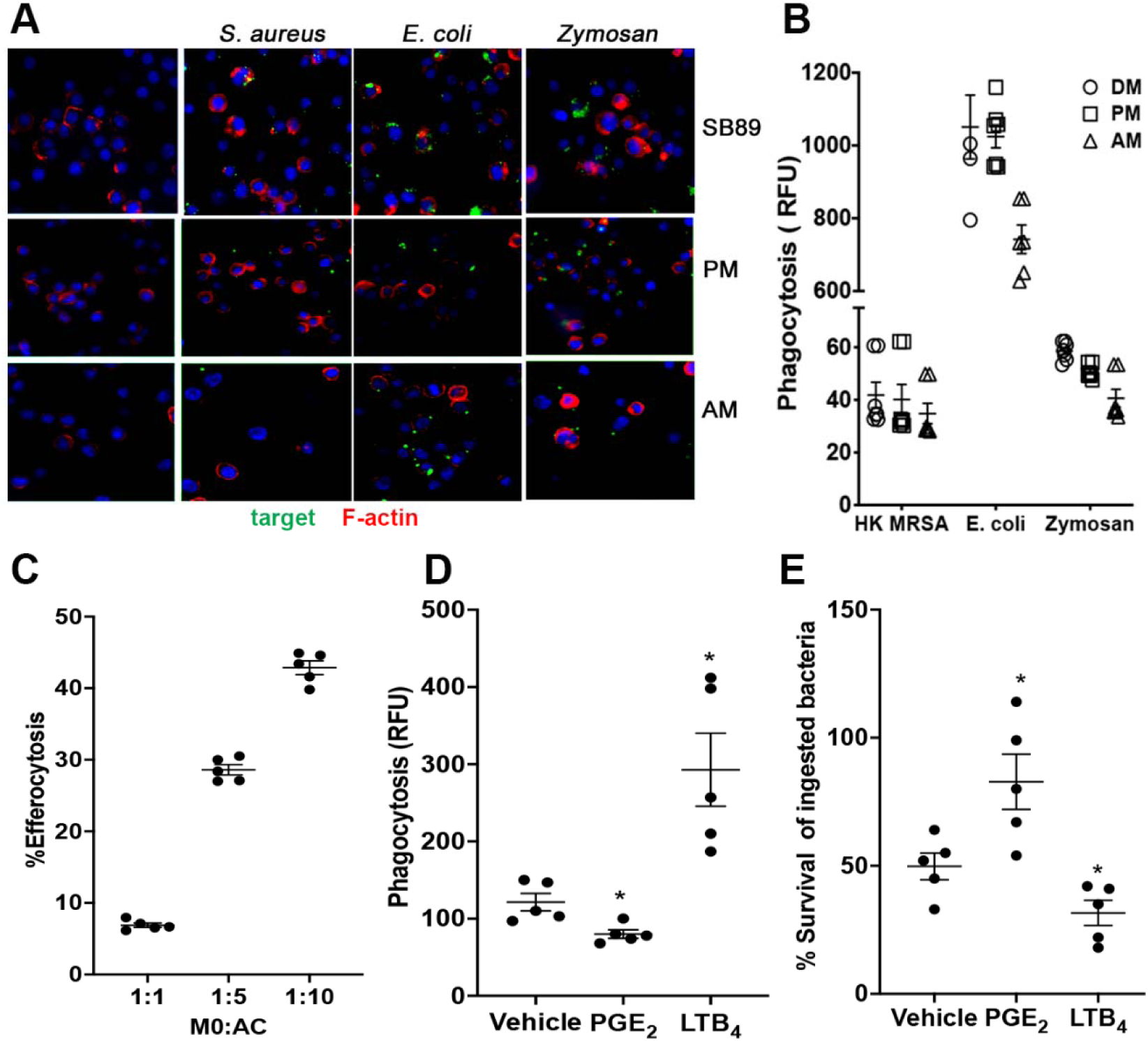
SB89 cells efficiently phagocytose and kill various targets. **A)** SB89 cells, primary peritoneal macrophages (PM), and alveolar macrophages (AM) were challenged with Phrodo-labeled *S. aureus*, *E. coli*, and zymosan for 60 min. Cells were then stained with TRITC-phalloidin and DAPI, and images were acquired as described in the Materials and Methods section. **C)** SB89 cells were challenged with the indicated CFSE-labeled apoptotic cells, and efferocytosis was quantified as described in the Materials and Methods section. **D and E)** SB89 cells were pre-treated with 1 μM PGE_2_ or LTB_4_ for 5 min, then challenged with GFP-expressing MRSA, and microbial ingestion **(D)** and killing **(E)** were quantified as described in the Materials and Methods section. Data represent mean ± SEM. \**p* < 0.05 vs vehicle control by Mann–Whitney test. n = at least 4/group.

### Production of inflammatory mediators by SB89 cells during MRSA infection

Dermal macrophages coordinate immune responses by secreting inflammatory mediators, including cytokines, chemokines, and bioactive lipid mediators^5, 6, 19^. To assess SB89 cells ’ ability to secrete and respond to MRSA infection, cells were challenged with MRSA or appropriate vehicle controls for 12 hours, followed by quantification of inflammatory mediators. Our results show that MRSA strongly stimulates the production of IL-1, IL-6, and IL-17A, as well as the neutrophil chemoattractant CXCL2 (Fig. 3A). In contrast, IL-10 levels decreased in MRSA-exposed SB89 cells (Fig. 3A). We also examined SB89 cells to generate bioactive lipid mediators during infection. Mass spectrometric measurement of released eicosanoids shows that SB89 cells release the prostanoids PGE_2_ and PGD_2_, as well as lipoxygenase products such as 12-HETE, 5-HETE, and LTB_4_ (Fig. 3B). Overall, these findings demonstrate that SB89 cells produce a broad range of inflammatory mediators (proteins and lipids) that significantly influence effector functions and inflammatory responses.

**Fig. 3.**
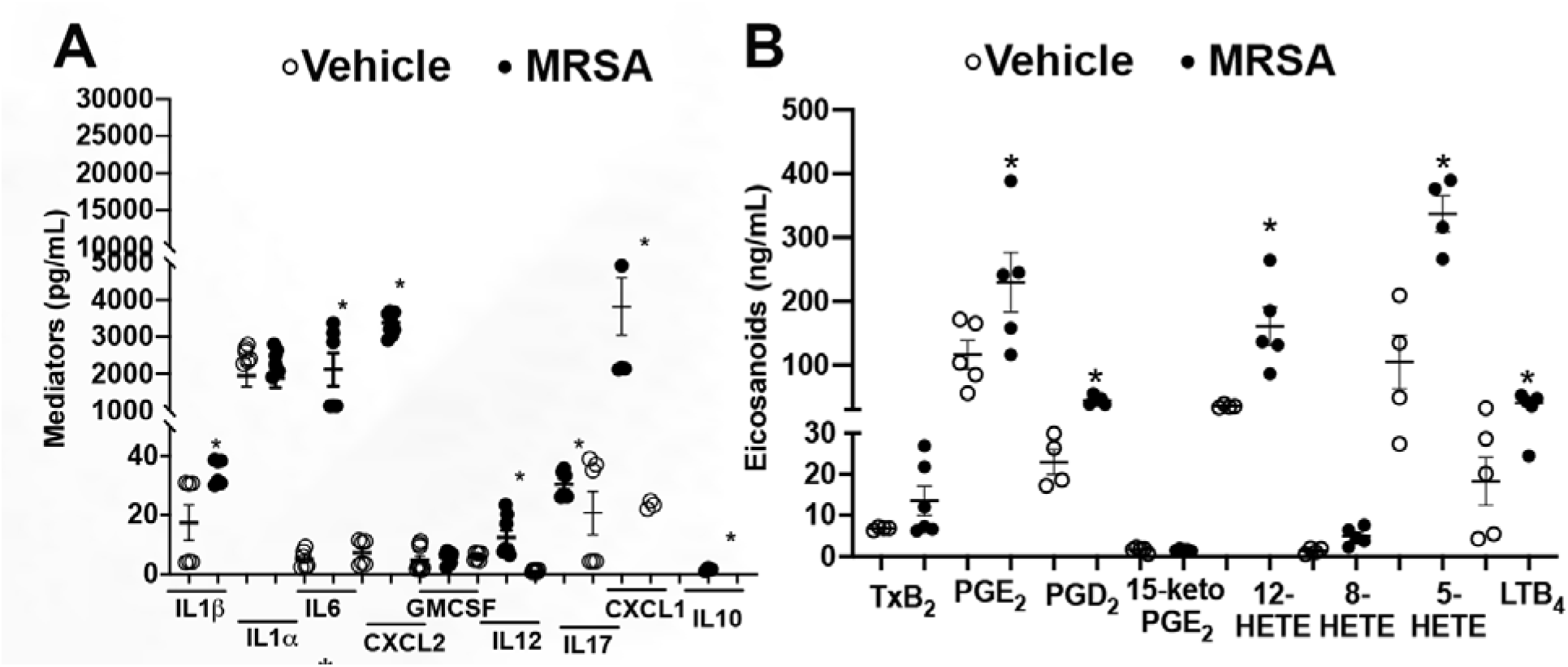
Production of pro- and anti-inflammatory mediators during MRSA infection. SB89 cells were challenged with MRSA (MOI of 10:1) for 18 h, and the production of **(A)** cytokines/chemokines and **(B)** eicosanoids was determined as described in the Material and Methods section. Data represent mean ± SEM. \**p* < 0.05 vs vehicle control by Mann–Whitney test. n = at least 4/group.

### SB89 cells are permissive to siRNA gene silencing and plasmid transfection

We next evaluated whether SB89 cells were permissive to gene silencing by RNA interference (RNAi). Given the role of microRNA-21 (miR-21) as a regulator of macrophage phenotype^20^, we have performed assays to silence miR-21 in SB89 cells. Figure 4A shows that miR-21 antagomir significantly decreased miR-21 expression in SB89 cells. The specificity of the silencing approach was confirmed by the absence of changes in miR125a expression by the miR21 antagomir (Fig. 4B). Consistent with our previous finding ^20^, inhibition of miR21 impaired LPS-induced nitric oxide and IL6 production by SB89 cells (Fig. 4C and D). In addition, SB89 cells were efficiently transfected with plasmid DNA encoding GFP, demonstrating their suitability for heterologous protein expression studies (Fig. 4E).

**Fig. 4.**
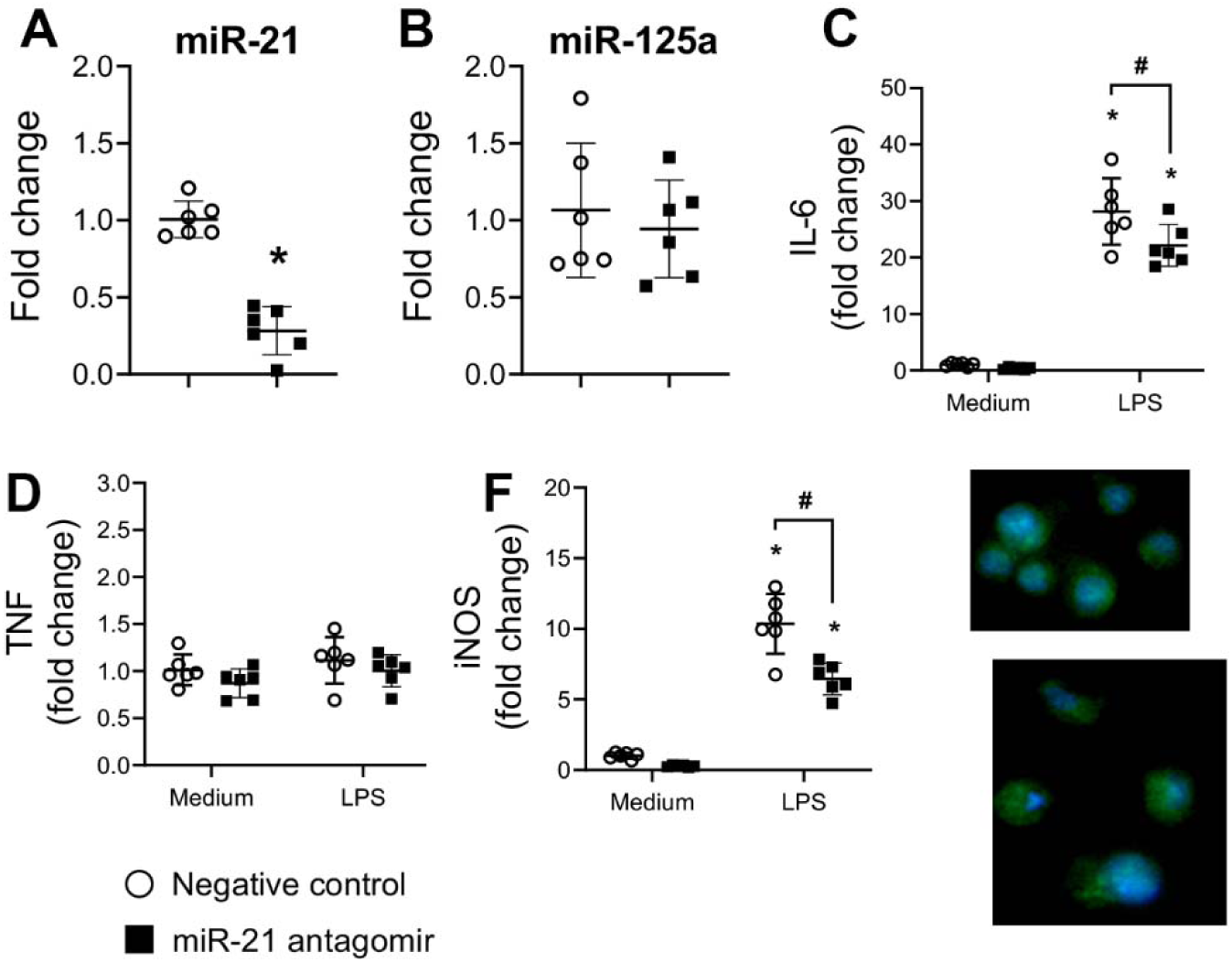
SB89 cells are permissive to transfection. **(A,B)** SB89 cells were transfected with miR21 antagomir or scrambled control as described in the Material and Methods section, and the expression levels of miR21 **(A)** and miR125a **(B)** were determined by qPCR. SB89 cells were transfected as in **A** and challenged with LPS for 24 h, followed by detection of IL-6 **(C)** and TNF **(D)** by ELISA. **(F)** SB89 cells were transfected and challenged with LPS as in C, and mRNA was isolated and Nos2 expression was detected by qPCR. **(E)** SB89 cells were transfected with a GFP-expressing plasmid and imaged as described in the methods, using citation 7. Data represent mean ± SEM. \**p* < 0.05 vs vehicle control by Mann–Whitney test. #P<0.05 VS scrambled control. n = at least 4/group.

### SB89 exhibits a distinct transcriptional profile compared with immortalized bone marrow derived macrophages (iBMDMs)

Next, we aimed to determine the basal transcriptional profile of SB89 cells using RNAseq analysis. To that end, we compared the transcriptomes of SB89 cells and immortalized BMDMs (iBMDMs). Differential gene expression analysis (Volcano analysis) identified 1740 downregulated and 2347 upregulated genes in SB89 cells relative to iBMDMs (Fig. 5A). Gene Ontology (GO) analysis of principal biological genes showed significant enrichment for genes involved in response to stimuli, defense response, and inflammatory processes (Fig. 5B and C), highlighting the strong immunological signature of SB89 cells. When we examined GO molecular function, we observed enrichment of genes involved in GPCR signaling, chemotaxis, and cell adhesion. Interestingly, genes associated with organ morphogenesis and neurobiological processes were also enriched (Fig. 5B and C). To further assess whether the transcriptional identity of SB89 cells differs from that of iBMDMs, we examined the transcription factors involved in the gene expression patterns described above. DMCL showed increased abundance of STAT3, ATF2, NFIC, JUN, HIF1A, and SMAD3. In iBMDMs, we observed increased expression of HOXC8, ETS1, RARG, and RELA. Together, these findings suggest that SB89 cells possess a transcriptional program distinct from that of iBMDMs, supporting the notion that they represent a unique skin-associated macrophage lineage.

**Fig. 5.**
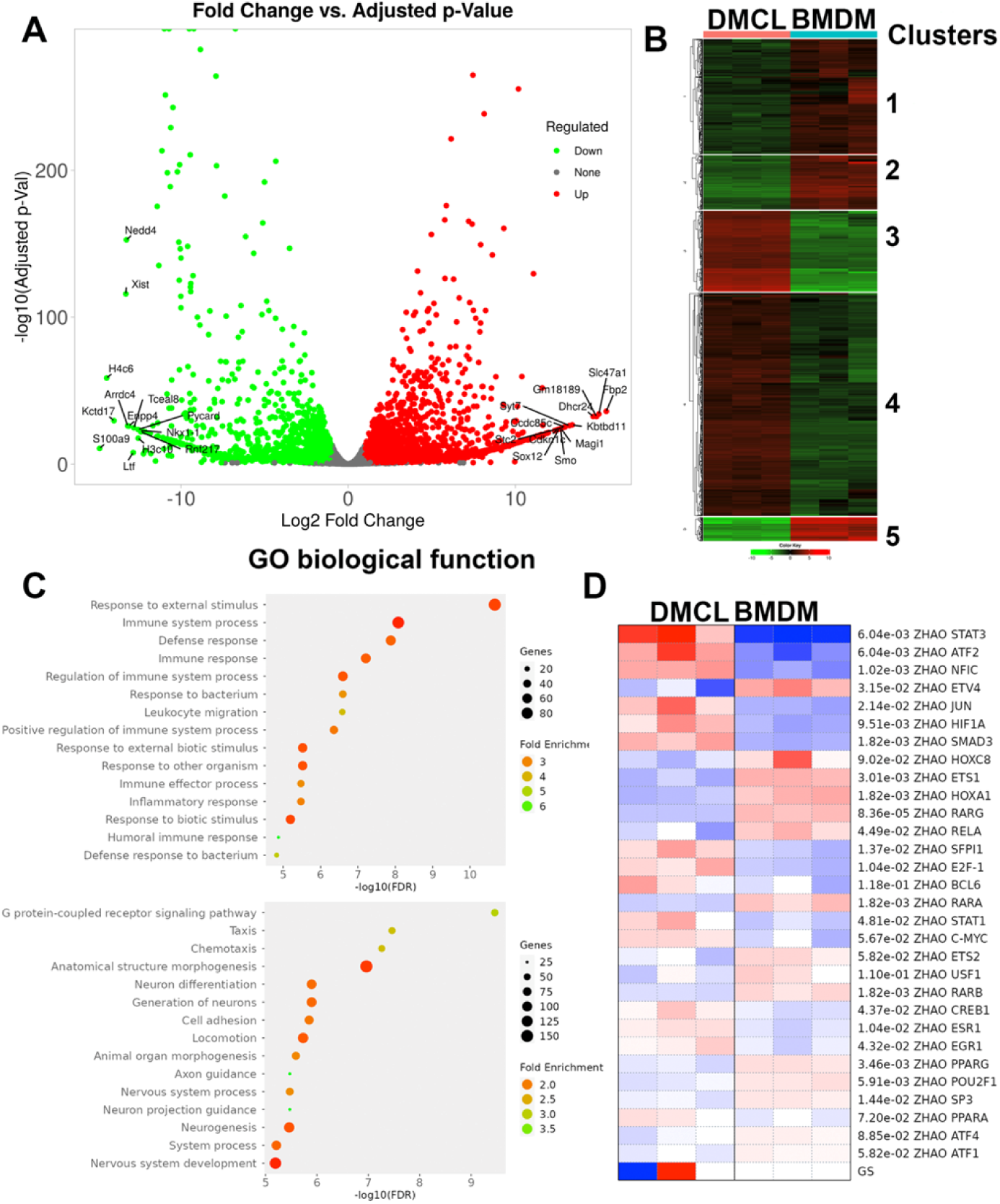
Differential gene expression analysis in SB89 cells and iBMDMs. SB89 cells and iBMDMs were collected, RNA was isolated, and samples were subjected to bulk RNAseq analysis. (A) Volcano plot (log-transformed) and (B) heat map showing significantly upregulated or downregulated genes in SB89 cells or iBMDMs. The listed genes correspond to the top 10 up- or downregulated genes. Gene analysis was performed using the iDEP Analysis Toolkit with FDR < 0.05. (C) Gene Ontology enrichment analysis of biological processes and molecular function. (D) Gene enrichment of transcription factors in SB89 cells and iBMDMs. RNAseq analyses were performed using iDEPP as described in the Material and Methods section. Data represent mean ± SEM from at least 3 independent experiments (RNA-seq).

## DISCUSSION

DMs play central roles in maintaining skin homeostasis and coordinating immune responses under both physiological and pathological conditions^7, 21, 22^. In healthy skin, these cells contribute to tissue repair, regulation of inflammation, maintenance of barrier integrity, and clearance of debris and pathogens^7, 21, 22^. In conditions such as chronic wounds, psoriasis, or skin cancers, these cells become dysregulated, leading to excessive inflammation, fibrosis, or tumor progression^7, 21, 22^. Despite its importance, studying DMs has been challenging due to their low abundance, difficulty in isolation, and phenotypic heterogeneity. Here, we offer an immortalized cell line derived from naive F4/80⁺ dermal macrophages that overcomes many of these limitations by eliminating the need for repeated animal testing and reducing batch variability associated with primary cells. Since SB89 cells were generated from non-activated skin macrophages, they are expected to retain tissue-specific transcriptional features that distinguish DMs from macrophages derived from other tissues, such as bone marrow or the peritoneal cavity.

Despite recent progress in understanding skin macrophage ontogeny, their heterogeneity and functions remain unclear^23, 24^. Addressing these questions is essential for developing targeted therapies that modulate macrophage biology and macrophage-driven inflammatory responses in the skin. Our newly developed cell line overcomes this limitation, providing a valuable tool for studying the unique properties and functions of DMs under both normal and disease conditions.

Skin phagocytes comprise heterogeneous populations whose phenotypes are shaped by local tissue microenvironmental and inflammatory signals. Skin-resident macrophages share several surface markers with CD11b^+^ dermal dendritic cells^25, 26, 27^. Tamoutounour et al. (2013)^3, 28^ identified six populations of CD11b^+^ cells in the dermis of C57BL/6 mice: one dendritic cell population, one monocyte-derived dermal macrophage (P1), two monocyte-derived dendritic cell populations (P2 and P3), and two resident dermal macrophage populations (P4 and P5).

Both P4 and P5 are positive for CD11b, CD64, and MerTK, and show low expression of CCR2 and Ly6C. However, they can be distinguished: P5 is positive for MHC class II and expresses low levels of CD11c, whereas P4 does not express either marker. Based on cell-surface markers (CD11b, CD64, CCR2, and MHC II), SB89 appears to be a P5 rather than a P4 resident dermal macrophage. Additionally, despite high F4/80 expression in SB89 cells, this marker alone is insufficient to discriminate CD11b^+^ cells in the dermis, as all six dermal populations show variable but significant F4/80 expression ^3, 28^.

Recent single-cell and bulk transcriptomic studies have further established that resident dermal macrophage subsets (P4/P5) differ from both circulating monocytes and other tissue macrophages in their gene expression profiles, turnover, and function. In that context, SB89 cells were derived from steady-state F4/80⁺/CD68⁺ dermal macrophages and retain a phenotypic profile (CD11b⁺ CD64⁺ CCR2^low^, MHC II⁺, F4/80, CD115, CD206 ^high^) that is highly consistent with the resident dermal macrophage subset described by Tamoutounour and colleagues that contributes to long-lived, self-renewing dermal macrophage pools. This alignment suggests that SB89 cells capture a bona fide resident, rather than a monocyte-derived inflammatory, dermal macrophage population, distinguishing them from transient CD14⁺ dermal macrophages and short-lived monocyte-derived dermal populations reported in other datasets.

Comparison of transcriptional signatures further highlights convergence and divergence between SB89 cells and primary dermal macrophages. RNA-seq analysis shows that SB89 cells upregulate genes involved in defense responses, inflammatory signaling, chemotaxis, cell adhesion, and GPCR signaling, reflecting core innate functions that are also enriched in dermal P4/P5 subsets in *in vivo* gene expression atlases. At the same time, SB89 cells show enrichment of pathways linked to organ morphogenesis and neurobiology-related functions, which parallels reports that dermal macrophages participate in nerve homeostasis and sensory function *in situ*, but also suggests that immortalization and prolonged culture in 2D conditions may exaggerate or rewire certain developmental and neuroimmune programs relative to freshly isolated primary cells.

Functionally, SB89 cells recapitulate key hallmarks of macrophage biology by efficiently phagocytosing bacteria (*S. aureus*, *E. coli*) and yeast-derived zymosan, as well as apoptotic cells. Interestingly, their phagocytic activity was comparable to or greater than that observed in PM and AM cell lines under similar culture conditions. In addition to internalization, SB89 cells efficiently kills MRSA, and its antimicrobial functions are dynamically regulated by lipid mediators: PGE₂ dampens, whereas LTB₄ enhances, phagocytosis and bacterial killing, mirroring known regulatory axes in primary macrophages^18^. Additionally, upon bacterial challenge, SB89 cells produce a broad spectrum of inflammatory cytokines and chemokines (including IL⍰1β, IL⍰6, IL⍰12p70, IL⍰17A, CXCL1, CXCL2) as well as eicosanoids (PGE₂, PGD₂, 5⍰HETE, 12⍰HETE, LTB₄). These findings indicate that SB89 cells retain the ability to integrate microbial and lipid-derived signals to shape the inflammatory milieu typical of infected or damaged skin.

*In vivo* studies have shown that primary dermal macrophages function as sessile scavengers that efficiently clear debris and pathogens but display limited migratory capacity and relatively modest T cell-priming potential compared with dermal dendritic cells^7, 22, 28, 29^. The strong phagocytic capacity, high expression of scavenger receptors, and dermal-resident surface phenotype of SB89 cells are consistent with this specialization, although its migratory and antigen-presentation functions remain to be directly compared with primary P4/P5 subsets. Together, these comparisons indicate that SB89 cells are a faithful surrogate for resident dermal macrophages, best suited to mechanistic studies focused on phagocytosis, inflammatory mediator production, and dermal imprinting, rather than on dynamic migratory behavior (since those endpoints were not investigated here) or complex tissue–tissue interactions that require intact skin architecture.

The J2 retroviral immortalization strategy used to generate SB89 cells has been successfully applied to bone marrow, splenic, fetal liver, microglial, and alveolar macrophages, producing cell lines that closely mirror their primary counterparts in polarization potential and basic macrophage functions. For example, J2-immortalized BMDMs maintain M1/M2 polarization profiles^30^, inflammasome activation^31^, and metabolic reprogramming that closely track those of primary BMDMs^12, 30^, thereby validating this approach for mechanistic studies of macrophage biology. In fact, SB89 cells share the following advantages: they exhibit typical macrophage morphology, express canonical macrophage markers, respond to TLR4 and bacterial stimuli with balanced pro- and anti-inflammatory responses, and support RNAi-based gene silencing and plasmid transfection. This makes SB89 cells a versatile source for mechanistic studies of signaling pathways, transcriptional regulation, and immunometabolism.

Importantly, SB89 cells also highlight the persistence of tissue-specific gene expression programs. In contrast to J2-immortalized BMDMs, whose transcriptional landscape largely overlaps with that of primary BMDMs, SB89 cells diverge markedly from iBMDMs, with thousands of differentially expressed genes and an overrepresentation of dermis-relevant signatures and transcription factors, including *Stat3, Hif1a, Jun, Atf2, Nfic*, and *Smad3*. This divergence underscores that J2-immortalized macrophage lines do not converge toward a common “generic” macrophage state; instead, they retain aspects of their tissue of origin. Therefore, SB89 cells complement, rather than replace, BMDM-derived J2 lines by providing a dermis-imprinted macrophage model. The comparative analyses above position SB89 cells as a dermis-committed macrophage line that bridges the gap between the scarcity of primary dermal macrophages and generic immortalized macrophage lines, thereby enabling detailed mechanistic studies grounded in tissue identity. SB89 cells should be particularly advantageous for dissecting pathways that integrate microbial cues, lipid mediators, and transcriptional regulators of dermal macrophage function, as well as for constructing reporter or CRISPR-edited derivatives to interrogate metabolism, signal transduction, and epigenetic regulation in a skin-relevant macrophage background. Nevertheless, as with any immortalized cell model, continued validation against primary dermal macrophages is critical. Future studies comparing SB89 cells with freshly isolated skin-resident macrophages using updated single-cell and bulk transcriptomic datasets, functional assays of migration and antigen presentation, and long-term passage studies will help define the boundaries of SB89 cells’ fidelity and avoid overextending conclusions beyond what this model can faithfully recapitulate in vivo.

SB89 provides a stable, dermis-imprinted macrophage model that preserves key resident features while enabling mechanistic, genetic, and high-throughput studies that are not feasible with primary dermal macrophages.

## MATERIAL AND METHODS

### Animals

C57BL/6 mice were purchased from Jackson Laboratory and bred at Vanderbilt University Medical Center (VUMC), an accredited animal facility. All procedures were approved by the Institutional Animal Care and Use Committee at VUMC (protocol #M1600215) and conducted in accordance with the Guide for the Care and Use of Laboratory Animals and the Public Health Service Policy on Humane Care and Use of Laboratory Animals.

### Isolation of skin resident macrophages

Mice were shaved, and residual fur was removed with a depilatory agent. Dorsal skin samples were removed and digested with collagenase IV (Merck KGaA, Darmstadt, Germany), as previously shown^32^. Flow cytometry was used both to sort dermal macrophages for transformation and to phenotype the clone that established the SB89 cell line. In both cases, cells were suspended in flow cytometry buffer (1% FBS in PBS) containing anti-CD16/32 to block nonspecific binding and incubated for 30 min at 4°C in the dark with the following fluorescence-conjugated antibodies for murine phagocyte surface markers (all from BD Biosciences): F4/80, CCR2, CD64, SIRPα, Cd11c, CD11b, CD115, Langerin, CCR7, MHCII, CX3CR1, CD206. Dermal macrophages were identified and sorted as F4/80^+^CD68^+^ cells. Samples were acquired on a FACSCanto II flow cytometer (BD Biosciences, Franklin Lakes, NJ). Data were analyzed using the FlowJo^TM^ software, version 10.5.3 (BD Biosciences, Ashland, OR).

To sort skin macrophages, the cell suspension was stained and sorted for macrophages (F4/80^+^ CD68^+^ double-positive cells) and macrophages were suspended in DMEM, supplemented with 10% heat inactivated fetal bovine serum (FBS), 2 mM l-glutamine, 100 units/ml penicillin, 100 μg/ml streptomycin, 25 mM HEPES and 2.5 × 10^5^ M 2-mercaptoethanol (all from Gibco Invitrogen, Grand Island, NY, USA) and seeded in plate dishes.

### J2 recombinant retrovirus preparation

The AMJ2-C11 cell line (an AM cell line) was used as the source of the J2 retrovirus^10, 31^. Supernatants from exponentially growing AMJ2-C11 cultures were collected, and cell debris was removed by centrifugation (3000 *x*g) for 15 min. The supernatants were filtered through a 0.45-μm membrane (Millipore, Bedford, MA), and 50% of the supernatant was incubated with 6 μg/ml polybrene (Sigma-Aldrich, St. Louis, MO, USA) and DMEM containing 10% FBS, L-glutamine (2 mM), penicillin (100 U/mL), and streptomycin (100 μg/mL).

### Immortalization of dermal macrophages

Primary DMs were isolated as described above and cultured in DMEM containing 100 ng/mL M-CSF for 24 h to stimulate proliferation. Nonadherent cells were washed off with warm medium 6–8 h later. Adherent DM monolayers were then infected with J2 virus supernatant mixed with DMEM-conditioned media and polybrene. After 24 h, extracellular virus was removed, and cells were subjected to two additional rounds of J2 infection at 24-h intervals under the same conditions to improve transfection efficiency. Clusters of rapidly proliferating cells were evident 14 days after initial infection. After 1 month, cells were grown in regular DMEM without M-CSF, and the proportion of virus-transformed cells progressively increased. Several rapidly growing DM cell lines were isolated in semisolid medium (0.96% methylcellulose) and subsequently recloned by limiting dilution. Individual clones were then isolated and characterized by staining for cell-surface markers associated with Langerhans’ cells, dermal, alveolar, and peritoneal macrophages.

### Peritoneal macrophage isolation

Resident peritoneal macrophages were isolated by PBS lavage from euthanized mice as previously described^33, 34^. Cells were centrifuged, and red blood cell lysis was performed if necessary. The cells were counted and plated for assays, and after at least 1 h of rest, the non-adherent cells were washed away with warm PBS.

### MRSA strains

The MRSA USA300 LAC strain was a gift from Dr. Bethany Moore (University of Michigan, Ann Arbor, Michigan, USA)^27^. The GFP-expressing USA300 LAC strain was a gift from William Nauseef (University of Iowa, Iowa City, Iowa)^29^. All MRSA stocks were stored at - 80 □C and cultured as previously described^30^.

### Phagocytosis assay

Bacterial phagocytosis was conducted as previously described^18^. SB89 cells, AM, and PM macrophages (2 × 10^5^ cells per well) were seeded into black-walled, clear-bottom 96-well plates (Cat #165305, Thermo Fisher Scientific, Rochester, NY, USA). Cells were cultured in DMEM and allowed to adhere for at least 1 hour. The medium was removed and replaced by complete DMEM containing FITC-labeled IHK-MRSA, pHrodo-labeled *E. coli,* or zymosan. GFP-MRSA was used at an MOI of 50:1 for 1 hour, and pHrodo-labeled bioparticles were added as recommended by the manufacturer￼. The plates were then washed with warm DMEM, and GFP fluorescence was measured. To assess intracellular fluorescence, extracellular GFP was quenched with 500 μg/mL trypan blue, and GFP levels were quantified using a fluorimeter plate reader at 488 nm, as we have shown. The trypan blue readings were used as a blank.

### Bacterial killing assay

Cells were pretreated with or without 1 μM LTB_4_ or PGE_2_ (Cayman Chemical Company, Ann Arbor, MI, USA) for 5 minutes prior to the addition of MRSA-GFP at a multiplicity of infection (MOI) of 50:1, as previously described^35^. Cells were incubated for 1 hour for phagocytosis, and both plates were washed with warm PBS to remove noninternalized bacteria. The PBS and treated samples were added to the killing plate and incubated for 2 hours for the killing assays. To measure the GFP fluorescence, extracellular fluorescence was quenched with 50 μL of 500 μg/mL trypan blue, and the intracellular GFP fluorescence was quantified by a plate-reader fluorimeter. Trypan blue alone served as the blank control. A reduction in GFP fluorescence in the killing plate relative to the phagocytosis plate indicated bacterial killing^36^.

### Generation of apoptotic cells

Thymus from C57BL/6 mice was harvested, and single-cell suspensions were generated by mechanical dissociation through a 70-um cell strainer. Cells were centrifuged, and red blood cell lysis was performed when necessary. Thymocytes were labeled with CFSE as previously described^37^ and cultured for 18 h to induce spontaneous apoptosis. Apoptotic cell frequency was determined by Annexin V and 7-AAD staining followed by flow cytometric analysis.

### Efferocytosis assay

DM were plated at a density of 1x10^6^ per well in a 6-well plate and cultured with apoptotic cells at different MOI (1:1, 1:5, and 1:10) for 2 h. After incubation, non-engulfed apoptotic cells were removed by washing, and the DM were harvested for flow cytometric analysis. Efferocytosis was quantified as a percentage of F4/80^+^CFSE^+^ cells^32, 37^.

### Multiplex bead array

SB89 cells (2 × 10^5^/well) were plated into two 96-well plates and maintained in complete DMEM (control group) or stimulated with 100 ng/mL ultrapure LPS (InvivoGen, San Diego, CA, USA) or MRSA (MOI of 50:1) for 24h. Total protein lysate was centrifuged at 10,000 *g* at 4 °C for 10 min to remove debris, and the supernatant was collected. Cell-free supernatant was collected and analyzed using the pro-inflammatory-focused 18-plex Discovery Assay from Eve Technologies (Calgary, AB) to detect cytokines and chemokines. Total protein content was determined by Bradford assay (Pierce) using bovine serum albumin as a standard. All concentrations were normalized to the total protein content of the corresponding tissue lysates and expressed as ng/mg protein.

### Detection of eicosanoids by mass spectrometry

DMCLs were exposed to MRSA (MOI of 50:1) for 6 hours, after which supernatants were collected. The Vanderbilt University Eicosanoid Core Laboratory processed the samples as previously described^38^. Eicosanoids were extracted with ice-cold methanol containing indomethacin and butylated hydroxytoluene (BHT). These samples were then analyzed using liquid chromatography-mass spectrometry (LC-MS). Eicosanoids were identified and quantified by mass and compared with known standards.

### Immunofluorescence

Macrophages were plated on 4- or 8-well glass chamber slides (Cat #154526 or #154534, Thermo Fisher Scientific) at a density of 1.25 × 10^5^ cells per well or 7.5 × 10^5^ cells per well, then infected with MRSA (MOI of 50:1). Phagocytosis was allowed to proceed for 30 minutes, after which the cells were washed three times with warmed DMEM. For subsequent time points, bacteria were removed after 30 minutes, and the slides were returned to the incubator until the specified time points. Cells were fixed with 4% methanol-free paraformaldehyde (Cat #47746, Cell Signaling Technology, Danvers, MA, USA) for 20 minutes. Following fixation, cells were washed, permeabilized in PBS containing 0.05% Triton X-100, and stained for F-actin using 0.165 µM AF 488 fluorescent phalloidin (ThermoFisher; #A12379) for 20 minutes in 1% BSA/PBS. After washing, slides were mounted with ProLong Gold Antifade Mountant containing DAPI.

### RNA isolation, reverse transcription, quantitative real-time PCR

Total RNA was isolated from the indicated cell lines using lysis buffer (RLT; Qiagen). The cDNA and real-time PCR were performed as previously published ^39^ using the gene-specific primers on a CFX96 Real-Time PCR Detection System (Bio-Rad). Primers were purchased from Integrated DNA Technologies (Coralville, IA, USA). Relative expression was calculated as described^39^.

### RNAseq and Functional Enrichment Analysis

SB89 or iBMDMs were cultured in DMEM and RNA was extracted using the RNeasy Mini Kit (Cat #74104, Qiagen Hilden, Germany) and then sent for processing at VANTAGE (Vanderbilt Technologies for Advanced Genomics), as previously described. Genes with log_2_FoldChange values greater than 2 or less than -2 and p-values under 0.05 were used to create log-normalized volcano plots. We performed Gene Set Enrichment Analysis (GSEA) on differentially expressed genes (DEGs) and genes with |log_2_FoldChange| ≥ 2, excluding low-abundance RNA reads. Pathway analysis was conducted using the KEGG database and the Gene Ontology (Biological Process, non-redundant) via the iDEEP gene analysis package, with an FDR < 0.05 as the screening threshold.

### Diff quick staining

SB89, AMJ2C11 cell lines and primary peritoneal cells were seeded in 8-well chambers and stained for Diff-Quik (Sigma-Aldrich). Images were visualized and acquired using the Nikon Eclipse Ci and Nikon Ds-Qi2 (Nikon, Tokyo, Japan).

### Statistical Analysis

The results are shown as the mean ± SEM and were analyzed using the Prism software, version 8.0.2 (GraphPad Software Inc., San Diego, CA, USA). Student’s *t*-test was used to compare two experimental groups, and one-way analysis of variance (ANOVA), followed by Bonferroni multiple comparisons tests, was used to compare three or more experimental groups. Values of *p* ≤ 0.05 were considered significant.

## ACKNOWLEDGMENTS

We would like to thank the Serezani laboratory for their input and support. This work was supported by NIH grants 1R01AI180777-01, R01DK122147 (to CHS). D.R was funded by 5T32AI112541-07. AS was funded by 5TL1TR002244-09. ASN was a recipient of a fellowship from Fundação de Amparo à Pesquisa do Estado de São Paulo (FAPESP, Grant # 2019/10839-4).

Genomic sequencing was performed in collaboration with the VANTAGE core services at Vanderbilt University, which is supported by the Vanderbilt Ingram Cancer Center (P30 CA068485) and the Vanderbilt Digestive Disease Research Center (P30 DK058404).

## AUTHOR CONTRIBUTION

CHS designed the research, analyzed the data, obtained funding, and wrote the article

SB designed research, performed experiments, and analyzed data

AJ, DR, and AS AB performed experiments and analyzed data

AS-N performed experiments, generated biological samples, and wrote the article

All authors read and approved the final manuscript.

